# *CHIQUITA1* maintains temporal transition between proliferation and differentiation in *Arabidopsis thaliana*

**DOI:** 10.1101/2021.11.24.469926

**Authors:** Flavia Bossi, Benjamin Jin, Elena Lazarus, Heather Cartwright, Yanniv Dorone, Seung Y. Rhee

## Abstract

Body size varies widely among species, populations, and individuals depending on the environment. Transitioning between proliferation and differentiation is a crucial determinant of final organ size, but how the timing of this transition is established and maintained remains unknown. Using cell proliferation markers and genetic analysis, we show that *CHIQUITA1* (*CHIQ1*) is required to maintain the timing of the transition from proliferation to differentiation in *Arabidopsis thaliana*. Combining kinematic and cell lineage tracking studies, we found that the number of actively dividing cells in *chiquita1-1* plants decreases prematurely compared to wild type plants, suggesting CHIQ1 maintains the proliferative capacity in dividing cells and ensures that cells divide a certain number of times. *CHIQ1* belongs to a plant-specific gene family of unknown molecular function and physically and genetically interacts with three close members of its family to control the timing of proliferation exit. Our work reveals the interdependency between cellular and organ-level processes underlying final organ size determination.

**Significance:** Timing of the transition between proliferation and differentiation is fundamental for determining the final size of organs and organisms. In agriculture, controlling organ and organism size can influence key agronomic traits such as yield and biomass. Dwarfism prevents lodging and was the trait responsible for the Green Revolution. Today, more sophisticated traits are needed for generating crops that are both resilient and sustainable. Revealing the molecular mechanisms that control the temporal transition between proliferation and differentiation will help unlock the potential of next-generation crops. Here, we report that *CHIQUITA 1* in *Arabidopsis thaliana* is needed to maintain the proper timing of the transition between proliferation and differentiation in leaves and roots.

## Introduction

Body size control is fundamental for growth and development and can impact reproductive fitness and ecosystem structure (1, 2). Understanding how plants control their body size has direct implications on agricultural productivity and resource utilization. While body size control is well established for insects and mammals (3, 4), much less is known for plants (5, 6).

Body size is driven by the rate and duration of organ growth (3), which determine the number of cell divisions and how large cells become. Differences in organ size among species are largely controlled by cell number rather than cell size (7, 8). The final cell number in an organ derives from the rate and duration of cell proliferation (9). In animals, cell death also contributes to the final cell number in an organ (8). The rate of cell proliferation within an organ depends on the number of dividing cells at any given time and the length of the cell cycle. Cell proliferation ends when all cells have exited the cell cycle. The genetic basis that controls proliferation exit during development is still unknown (10).

Three cell-intrinsic mechanisms of proliferation exit (i.e. intracellular mechanisms that stops proliferation, (8)) have been proposed to control the timing of the transition from proliferation to differentiation: 1) a timer mechanism (cells divide for a fixed period of time); 2) a counter mechanism (cells undergo a fixed number of divisions); and 3) a sizer mechanism (an optimal threshold cell size is required for cell cycle progression). Exit from proliferation is also controlled by systemic mechanisms at the organ or organism level, including long-distance signaling and cell-to-cell contact interactions (3). In animals, organs and organisms have evolved different strategies to control when cells exit proliferation (4, 11–18), indicating that no universal mechanism controls the timing of proliferation exit. For example, a timer composed of at least two cyclin-dependent kinase inhibitors controls how many times an oligodendrocyte precursor (11) and a myocyte divide (12). A cell sizer mechanism controls the number of cell divisions in *Xenopus laevis* embryos before the onset of the zygotic genome activation (ZGA) (16). Before ZGA, cells divide without growing, which reduces cytoplasmic volume and increases DNA concentration and DNA:cytoplasm ratio. When individual cells in *X. laevis* embryos reach a threshold size and DNA:cytoplasm ratio, ZGA is triggered, leading to the onset of germ-layer specification and cell differentiation (16). In plants, while a cell cycle counter mechanism has been proposed for developing energy sink organs such as flowers and roots (19, 20), it is unknown which mechanisms operate to control proliferation exit of the major energy source organ, the leaf.

We previously used a bioinformatic pipeline to find novel transcriptional regulators and identified *CHIQUITA1* (*CHIQ1*), a gene of unknown function involved in organ size control in *Arabidopsis thaliana* (21). We found that CHIQ1 physically interacts with other CHIQ-like proteins (21) and the Polycomb repressive complex 2 (PRC2), a transcriptional repressor complex, via another member of the CHIQ family (21). More recently, CHIQ1 (renamed CONSTITUTIVELY STRESSED 1 (COST1)) was implicated in the regulation of autophagy in response to drought stress (22). However, CHIQ1/COST1’s role during growth is independent of autophagy (22). Here, we show that CHIQ1 is a positive regulator of body size by measuring organ growth at the cellular, organ, and organismal levels using genetic and chemical perturbations combined with live 4D imaging. Genetic interaction studies indicated that *CHIQ1* works with three other previously uncharacterized *CHIQ*-like genes (*CHIQUITA 1-LIKE 4* (*CHIQL4*), *CHIQUITA 1-LIKE* 5 (*CHIQL5*), and *CHIQUITA 1-LIKE 6* (*CHIQL6*)) to control the timing of transition from proliferation to differentiation and final cell number in leaves. Kinematic and cell lineage tracking analyses indicate that CHIQ1 maintains proliferative capacity in dividing cells and imply that CHIQ1 ensures that cells divide a certain number of times before entering differentiation. Finally, we show that four members of the CHIQ family act together as positive regulators of cell proliferation during organ growth and are required for proper timing of cell cycle exit in leaves.

## Results

### CHIQ1 is a positive regulator of vegetative and reproductive organ size

*CHIQ1* encodes a protein of unknown function, which is required for *Arabidopsis thaliana* to reach full size at maturity (21). Previously we showed that the lines carrying the null allele *chiq1-1* had smaller leaves and stems than wild type plants (21). To determine whether other major organs, such as flowers, siliques, and roots, exhibited similar size reduction, we examined them at various stages of development. At maturity, these organs were significantly smaller than those in the wild type (Fig. 1). However, during the first week of growth following germination, where most growth occurs by cell division, *chiq1-1* organs were similar or even bigger than wild type organs (Fig. 2, Fig. S1). Primary roots of *chiq1-1* seedlings were ~10% longer than those of wild type plants 3-5 days after sowing. However, *chiq1-1* roots became ~20-40% shorter than wild type roots as the seedlings aged (Fig. 2A, Fig. 1B). During rosette growth, *chiq1-1* leaves ranged from being larger than the wild type (1^st^ and 2^nd^ leaves) to being similar to the wild type (3^rd^-7^th^ leaves), but all *chiq1-1* leaves became smaller than wild type leaves as the plants aged (Fig. 2C-D, Fig. S1). Introducing wild type *CHIQ1* into the mutant background complemented the organ size phenotypes observed in the *chiq1-1* null mutant (Fig. 1, Fig. 2).

**Figure 1.**
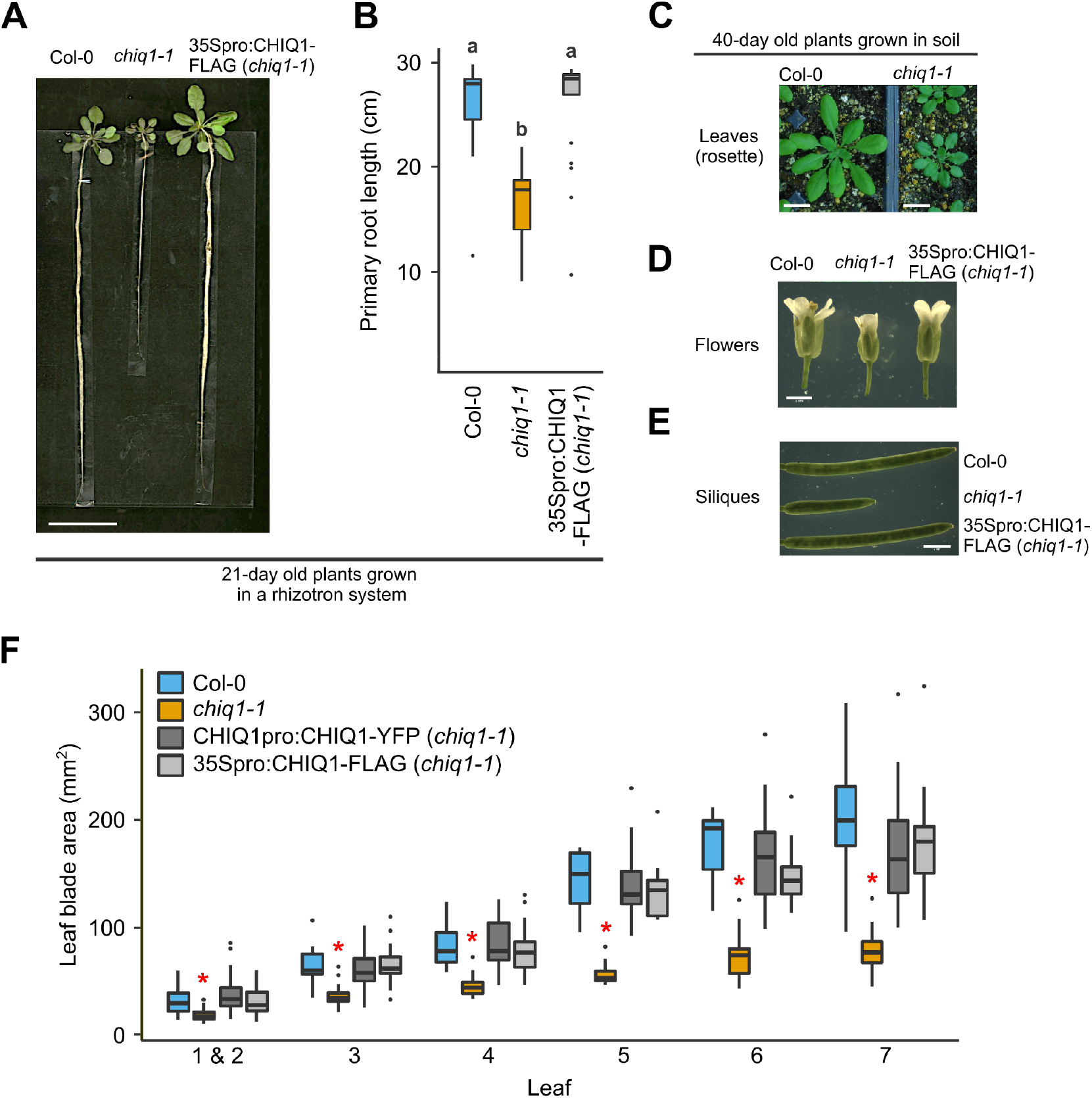
*chiq1-1* mature organs are smaller than wild type. A) 21 day-old plants grown in soil using a rhizotron system (Rellan-Alvarez et al., 2015) (wild type (left), *chiq1-1* (middle), and a complemented line (*35Spro:CHIQ1-FLAG*, right)). Scale bar = 5 cm. B) Length of roots of wild type (Col-0, blue), *chiq1-1* (orange) and one complemented line (*35Spro:CHIQ1-FLAG*, gray) grown in soil for 21 days using a rhizotron system (Rellan-Alvarez et al., 2015) (n= 20-21 per genotype; N=3). Letters represent significantly different groups (p-value < 0.05) as determined by Kruskal-Wallis test followed by pairwise comparison using the Wilcoxon test. C) Rosette leaves from 40 day-old wild type (left) and *chiq1-1* (right) plants. Scale bar = 2 cm. D) Flowers from wild type (left), *chiq1-1* (middle), and a complemented line (*35Spro:CHIQ1-FLAG*, right). Scale bar = 1 mm. E) Siliques from wild type (top), *chiq1-1* (middle), and a complemented line (*35Spro:CHIQ1-FLAG*, bottom). Scale bar = 2 mm. F) Leaf blade area (box and whisker plot) of mature leaves from soil-grown wild type (blue), *chiq1-1* (orange), and two complemented lines (*CHIQ1pro:CHIQ1-YFP* line, left gray bar and *35Spro:CHIQ1-FLAG*, right gray bar) (n= 6-71 per genotype per leaf; N=2). The blade area of each leaf was compared among the four genotypes using Kruskal-Wallis test followed by pairwise comparison using the Wilcoxon test. The red asterisk indicates statistical significance in the *chiq1-1* leaf compared to the other three genotypes (p-value < 0.05). B,F) Graphs were generated using the ggplot2 package in R (49).

**Figure 2.**
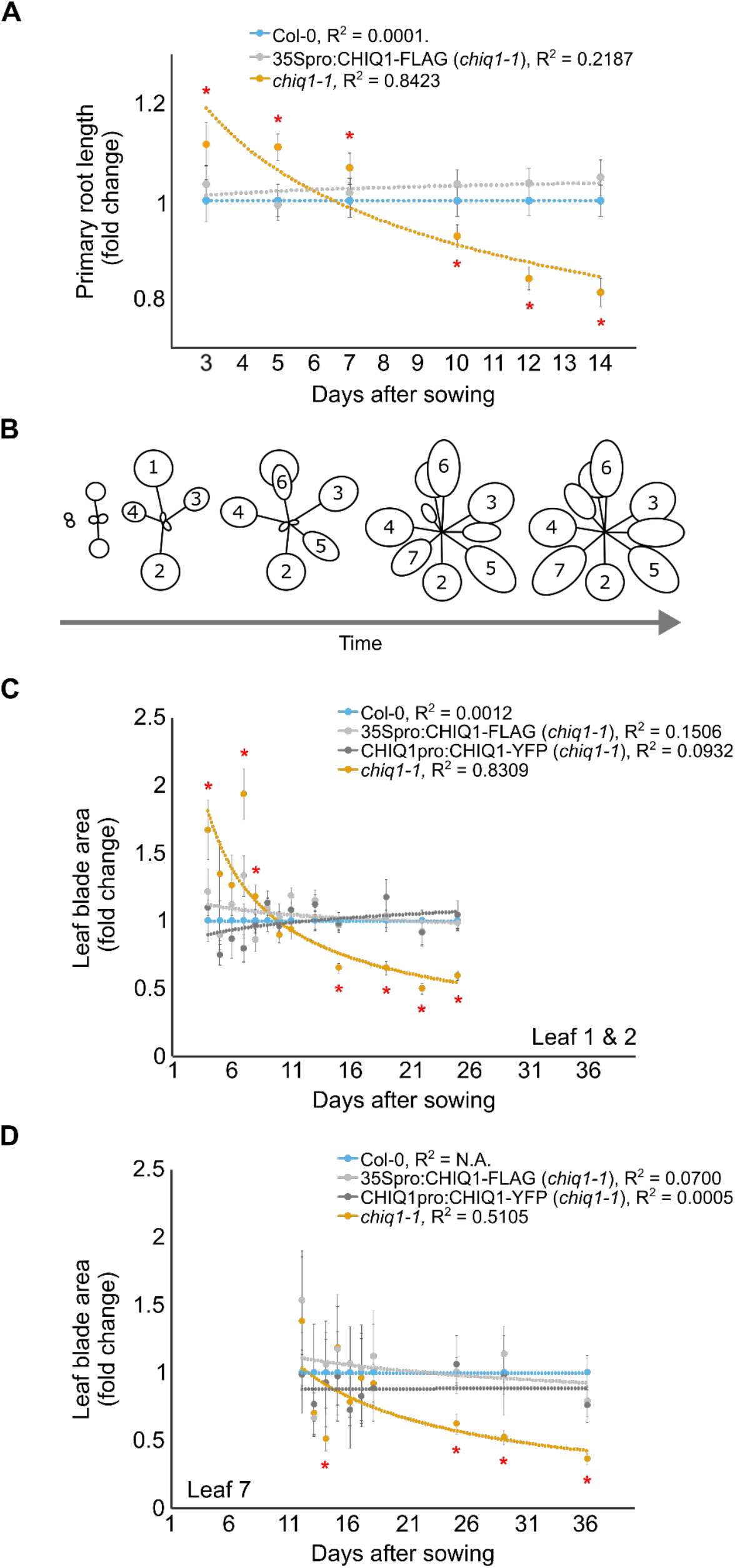
*chiq1-1* young organs are of similar size or slightly bigger than in wild type. A) Scatter plot of normalized primary root length against the wild type value of the same age in wild type (Col-0, blue), *chiq1-1* (orange), and one complemented line (35Spro:CHIQ1-FLAG, gray) grown in media plates from 3 to 14 days after sowing (n = 92-131 per genotype per time point; N = 3). Error bar represents 95% confidence interval. The regression line corresponds to a power regression. All genotypes were compared at each time point using Kruskal-Wallis test followed by pairwise comparison using the Wilcoxon test (p-value < 0.05). Red asterisks represent a significant difference between *chiq1-1* and the other two genotypes. B) Schematic representation of leaf growth in Arabidopsis rosette over time. C-D) Scatter plot of normalized leaf blade area against the wild type value of the same age of the first pair of leaves (C) and the 7^th^ (D) leaf in soil-grown plants ranging in age from 4 days to 36 days from wild type (blue), *chiq1-1* (orange), and two complemented lines (CHIQ1pro:CHIQ1-YFP line, dark gray and 35Spro:CHIQ1-FLAG, light gray) (n = 18-108 per genotype; N = 1-2 (1^st^ leaf) and n = 7-21 per genotype; N=1 (7^th^ leaf)). Error bar represents 95% confidence interval. The regression line corresponds to a power regression. All genotypes were compared at each time point using Kruskal-Wallis test followed by pairwise comparison using the Wilcoxon test (p-value < 0.05). Red asterisks represent a significant difference between *chiq1-1* and the other three genotypes. Graphs were generated using the ggplot2 package in R (49).

To determine whether the reduced organ size derives from a reduction in cell size or cell number, we first measured these parameters in plants at maturity. Fully expanded leaves in *chiq1-1* had both fewer and smaller cells (Fig. 3 A, B, E and G, Fig. S2). The same was observed in petals (Fig. 3 C, D, F and H). Since smaller cells could result from defects in endoreduplication (23, 24), we asked whether endoreduplication was affected in *chiq1-1* plants. The majority (59%) of *chiq1-1* cells were diploid (2C) in mature leaves compared to 31% in wild type cells (Fig. 3I). This indicates that the majority of *chiq1-1* leaf cells did not enter endoreduplication after exiting the mitotic cell cycle, which could explain the reduced cell size observed in *chiq1-1* leaves at maturity. While most *chiq1-1* cells skip endoreduplication, suggesting the transition from mitosis into endoreduplication is disrupted, pavement cells display the typical jigsaw puzzle shape in mature leaves, which suggests that terminal differentiation is not compromised in *chiq1-1*.

**Figure 3.**
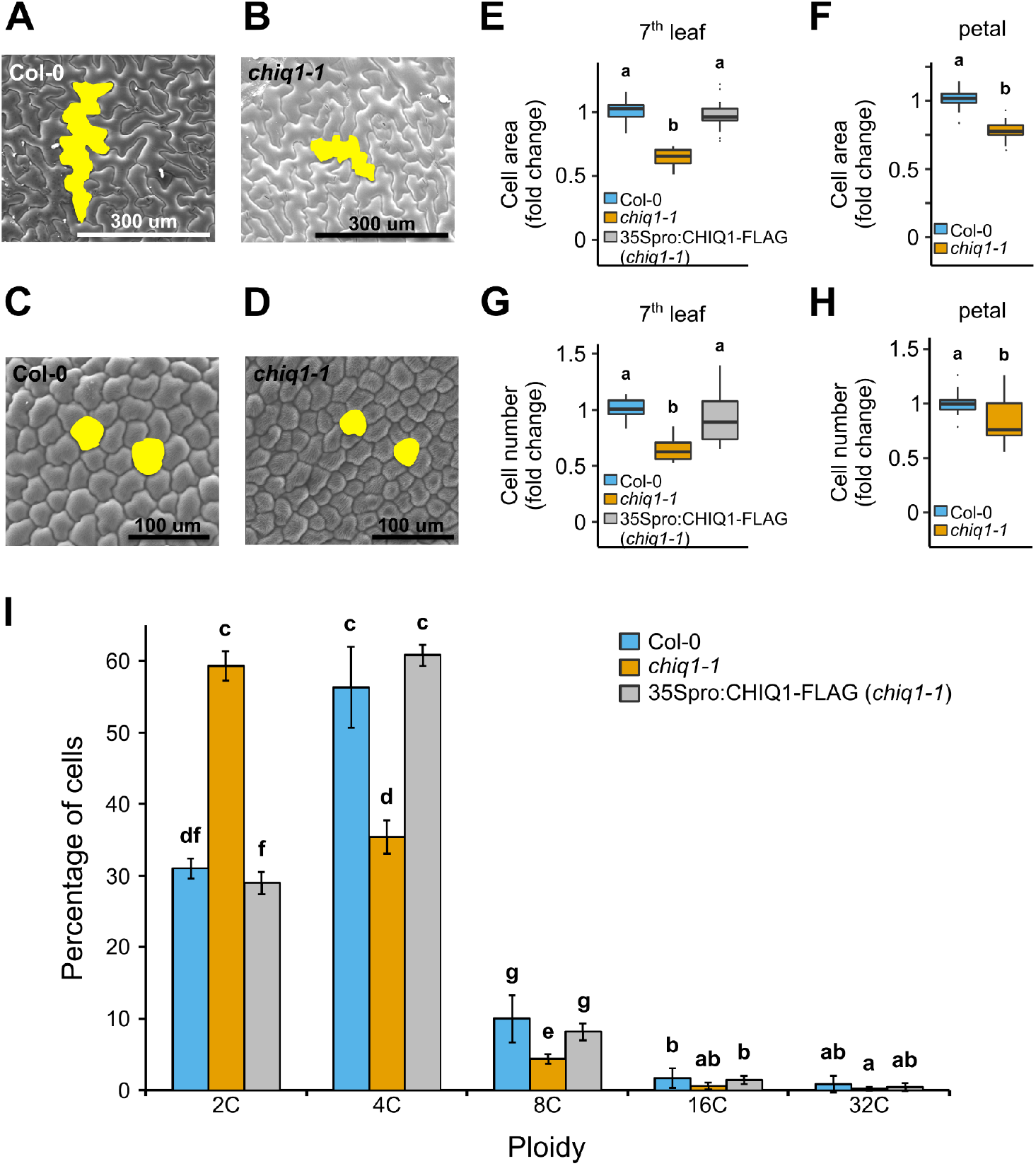
*chiq1-1* mature organs have fewer and smaller cells and cells undergo less endoreduplication in *chiq1-1* leaves. A-D) Scanning electron microscopy (SEM) images of the adaxial epidermis of the 7th leaf (A, B) and petal (C, D) from 51 day-old wild type (A, C) and *chiq1-1* (B, C) plants. E,G) Normalized average cell size (E) and total cell number (G) in the abaxial epidermis of the 7^th^ leaf against the wild type value of the same experiment from 36 day-old wild type (Col-0, blue), *chiq1-1* (orange) and a complemented line (35Spro:CHIQ1-FLAG, gray) (n= 12-24 per genotype; N=4). F, H) Normalized average cell size (F) and total cell number (H) of petal conical cells from wild type (Col-0, blue) and *chiq1-1* (orange) open flowers (n= 13-14 per genotype; N=3). Letters in E-H represent significantly different groups (p-value < 0.05) as determined by two-way analysis of variance followed by post hoc Tukey’s test. I) Percentage of cells with each ploidy level in mature leaves from wild type (Col-0, blue), *chiq1-1* (orange) and a complemented line (35Spro:CHIQ1-FLAG, gray) (n= 4 per genotype; N=2). Error bar represents 95% confidence interval. Letters represent significantly different groups (p-value < 0.05) as determined by Kruskal-Wallis test followed by pairwise comparison using the Wilcoxon test. Graphs were generated using the ggplot2 package in R (49).

Taken together, our data show that *CHIQ1* affects final organ size by modulating cell proliferation and expansion. We next investigated the role of *CHIQ1* during cell proliferation and the transition from cell proliferation into cell expansion.

### *CHIQ1* maintains cell number and cell size during development

To determine what led to the late onset dwarfism in *chiq1-1*, we compared how cell proliferation and expansion changed over time between *chiq1-1* and wild type plants by employing a kinematic analysis on the first pair to leaves from 4 to 25 days after sowing. Total cell number was similar in both genotypes from day 4 to day 15 after sowing and became significantly lower at days 19 and 25 after sowing in *chiq1-1* leaves compared to the wild type (Fig. 4A). The number of cells in *chiq1-1*’s root apical meristem (RAM) decreased similarly over time (Fig. S3). The stomatal index (SI)—which represents the proportion of guard cells (specialized cells involved in gas exchange) among all cells in the leaf epidermis—was similar in both genotypes (Fig. S4). This indicates *CHIQ1* does not affect the pattern of cell type differentiation in the epidermis.

**Figure 4.**
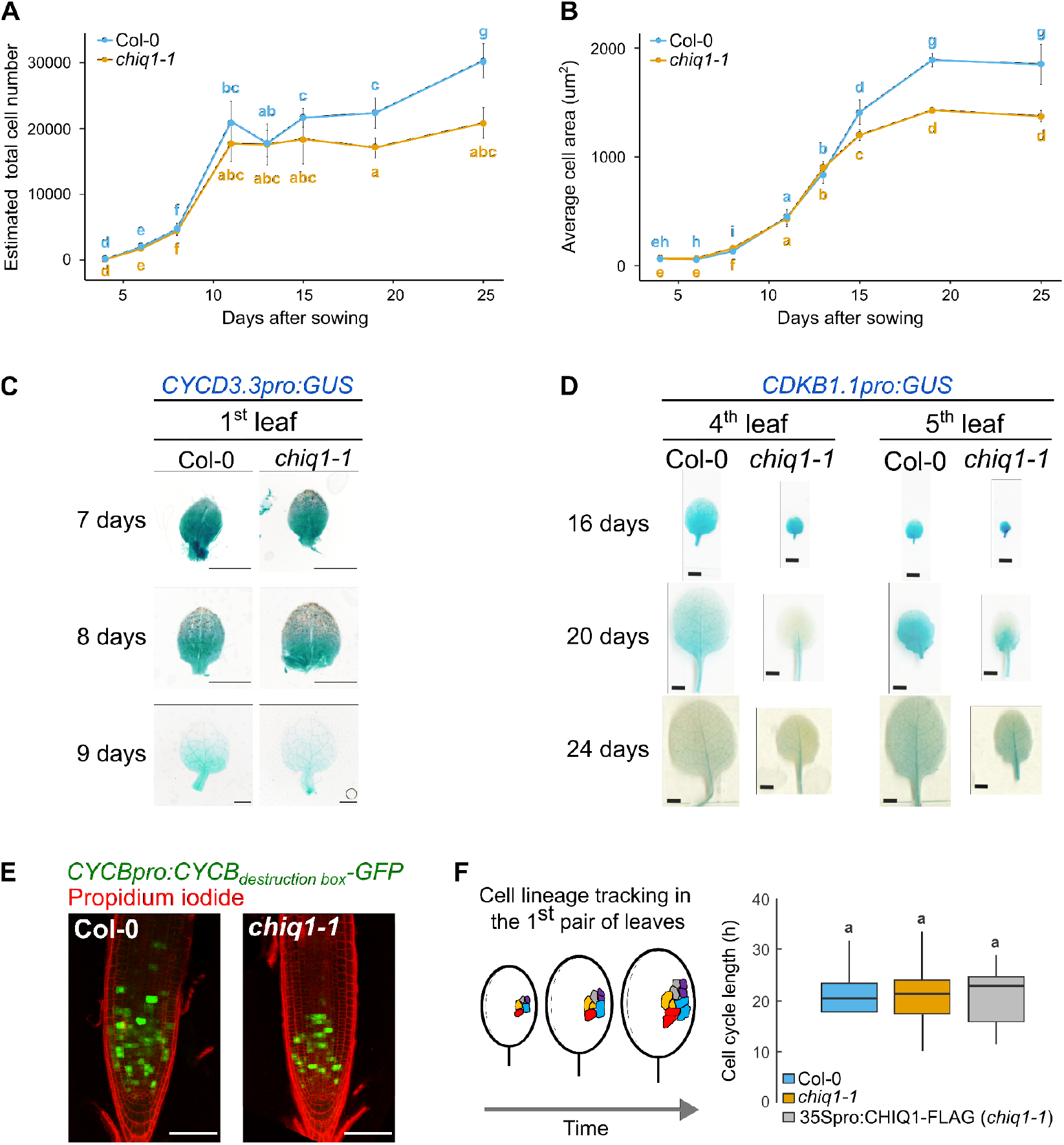
Fewer cells divide in *chiq1-1* and *chiq1-1* cells exit proliferation prematurely in growing organs. A-B) Kinematic studies in the first pair of true leaves from wild type (Col-0, blue) and *chiq1-1* (orange) seedlings grown on soil. Estimated total cell number (A) and average cell area (B) in the abaxial epidermis from wild type (Col-0, blue) and *chiq1-1* (orange) seedlings at 4, 6, 8, 11, 13, 15, 19 and 25 days after sowing (n = 5-30 per genotype per time point; N = 1-3). Letters represent significantly different groups (p-value < 0.05) as determined by Kruskal-Wallis test followed by pairwise comparison using the Wilcoxon test. C) Expression of *CYCD3.3pro:GUS* in wild type (Col-0, left) and *chiq1-1* (right) first true leaves grown for 7, 8, and 9 days. Every image is representative of two independent lines. A total of ten leaves were analyzed per genotype per experiment. Scale bar = 0.5mm. D) Expression of *CDKB1.1pro:GUS* in the 4^th^ and 5^th^ leaves of wild type (Col-0, left) and *chiq1-1* (right) plants at days 16, 20, and 24 after sowing. Every leaf is a representative image of two independent experiments and four leaves were analyzed per genotype per experiment. Scale bar = 2 mm. E) Expression of the cell cycle marker *CYCBpro:CYCB_destruction box_*-GFP in the root apical meristem of 5 day-old wild type (left panel) and *chiq1-1* (right panel) seedlings stained with propidium iodide. Scale bar = 100um. F) Left panel: schematic representation of the experimental design and cell lineage tracking analysis in leaves. The first pair of leaves was imaged daily from day 6 to 8 after sowing. Images were taken from the whole leaf and cells in the middle region of the leaf blade were traced and followed for 2 days (more details in text). In this representation, cells with the same color belong to the same lineage. Right panel: Cell cycle length from day 6 to day 8 in leaves from wild type (Col-0, blue), *chiq1-1* (orange), and a complemented line (35Spro:CHIQ1-FLAG, gray) seedlings (n = 17-35 per genotype; N = 3-7). Letters represent significantly different groups (p-value < 0.05) as determined by two-way analysis of variance followed by post hoc Tukey’s test. Graphs were generated using the ggplot2 package in R (49).

Difference in cell size between wild type and *chiq1-1* leaves showed similar dynamic patterns over time as the cell number. The average cell area was greater in *chiq1-1* at 4-8 days after sowing (Fig. 4B, Fig. S5A), which could explain the temporary increase in leaf size at early developmental time points (Fig. 2C). Later in development, the average cell area was similar in both genotypes at 11-13 days after sowing, and became smaller in *chiq1-1* seedlings starting at day 15 after sowing (Fig. 4B). Because the leaf epidermis is composed of cells that are orders of magnitude different in size depending on cell type (guard vs. pavement) and differentiation state (dividing vs. differentiated), we wondered whether examining only one cell type would reveal further insight. Therefore, a more detailed analysis of the cell area distribution of epidermis cells excluding guard cells was performed. We found that the proportion of small cells—which likely represent dividing cells based on previous studies (25, 26)—was smaller in 8 day-old *chiq1-1* leaves compared to wild type leaves (Fig. S5B). This suggests that the proportion of dividing cells is smaller in 8 day-old *chiq1-1* leaves while the proportion of expanding cells is larger than the wild type, which would explain the ephemerally larger leaves in 8-day old *chiq1-1* plants compared to the wild type. These data suggested that cell proliferation is decreasing faster in *chiq1-1* organs and that cells may be transitioning prematurely into a differentiated state.

### *CHIQ1* delays exit from cell proliferation

To test the hypothesis that CHIQ1 may be involved in controlling the exit from cell proliferation, we followed expression patterns of cell cycle markers over time in actively growing leaves and roots. For cell cycle markers, we used *CYCLIN D3.3 (CYCD3.3), CYCLIN B (CYCB1.1)*, and *CYCLIN-DEPENDENT KINASE B1.1 (CDKB1.1)*, which encode components of the cell cycle machinery expressed during cell proliferation (27–29). We found that the domain of expression of *CYCD3.3* and *CDKB1.1* became more restricted earlier in *chiq1-1* leaves (Fig. 4 C and D). Similarly, fewer cells expressed *CYCB1.1* in the RAM of 5 day-old *chiq1-1* seedlings (Fig. 4E). Together, these results suggest that fewer cells were dividing in *chiq1-1* organs, supporting the hypothesis of earlier exit from proliferation. Alternatively, since *CYCB1.1* expresses in late-G2/M, *chiq1-1* cells may be undergoing a longer cell cycle due to being arrested in the cell cycle phases G1 or S.

### Cells divide fewer times before exiting proliferation in *chiq1-1* leaves

To distinguish between fewer cells dividing or a longer cell cycle in *chiq1-1* leaves, individual cells expressing the fluorescent epidermis-specific plasma membrane marker RCI2A (30) were tracked from epidermis images of intact first true leaves from 6-day-old seedlings every 24 hours for 2 days (Fig. 4F). Cell cycle length was estimated from the proliferation rate, which was calculated by counting the cells that divided during the course of the experiment and their progeny. Cells that did not divide during the experiment were not included in the calculation of proliferation rate. We found that the cell cycle length was not significantly different between wild type, *chiq1-1*, and the complemented line 35Spro:CHIQ1-FLAG (Fig. 4F), indicating that the rate of cell cycle progression is similar in all genotypes. To further confirm that CHIQ1 does not affect cell cycle progression, we treated plants with hydroxyurea (HU), which inhibits the enzyme ribonucleotide reductase, reduces the amount of dNTPs available for DNA synthesis, and delays entry into mitosis (31). HU increased the cell cycle length in wild type and *chiq1-1* leaves indistinguishably (Fig. S6). All these data together indicate that *CHIQ1* does not compromise the rate of cell cycle progression and, more importantly, that the decrease in the cell proliferation rate at the leaf level is not due to a longer cell cycle but to a defect in the timing of proliferation exit. This premature proliferation exit decreases the population of dividing cells in *chiq1-1* organs leading to smaller mature organs with fewer cells in *chiq1-1* than in wild type plants.

### CHIQ1 works with other CHIQ-like proteins to control the timing of proliferation exit

CHIQ1 physically interacts with other CHIQ-like proteins (21). To test whether they work together genetically and understand the role of CHIQ proteins during organ growth, we generated a quadruple mutant called *chiq-quad*, which lacks CHIQ1, CHIQL4, CHIQL5 and CHIQL6, and analyzed leaf area, cell number, and cell size during development and at maturity. Mature leaf area in both the single and quadruple mutants was smaller than that in the wild type (Fig. 5A), but not different between *chiq1-1* and *chiq-quad* (Fig. 5A). However, cell number in *chiq-quad* leaves was further reduced compared to that in *chiq1-1* leaves (Fig. 5B, Fig. S7 A and B), which is consistent with these CHIQ1-like proteins participating in cell proliferation. Interestingly, the average cell area in *chiq-quad* leaves was greater than that in *chiq1-1* leaves and similar to the wild type (Fig. 5C, Fig. S7 C and D), which explains why the leaf area of *chiq-quad* and *chiq1-1* plants was indistinguishable despite having fewer cells. This supports a compensatory mechanism that could have been triggered as *chiq-quad* failed to reach a certain cell number threshold (32). In the growing leaf, a larger proportion of cells exited proliferation earlier in *chiq-quad* compared to *chiq1-1* and wild type as seen by the earlier decrease in cell number (Fig. 5E). In addition, cell size (Fig. 5F) and stomatal index (Fig. 5G) increased earlier in *chiq-quad* compared to *chiq1-1*, suggesting that cells in *chiq-quad* undergo cell differentiation even earlier than *chiq1-1* mutants. These data indicate that CHIQ proteins may be part of a complex that regulates the duration of cell proliferation at the organ level and the timing of cell differentiation onset.

**Figure 5.**
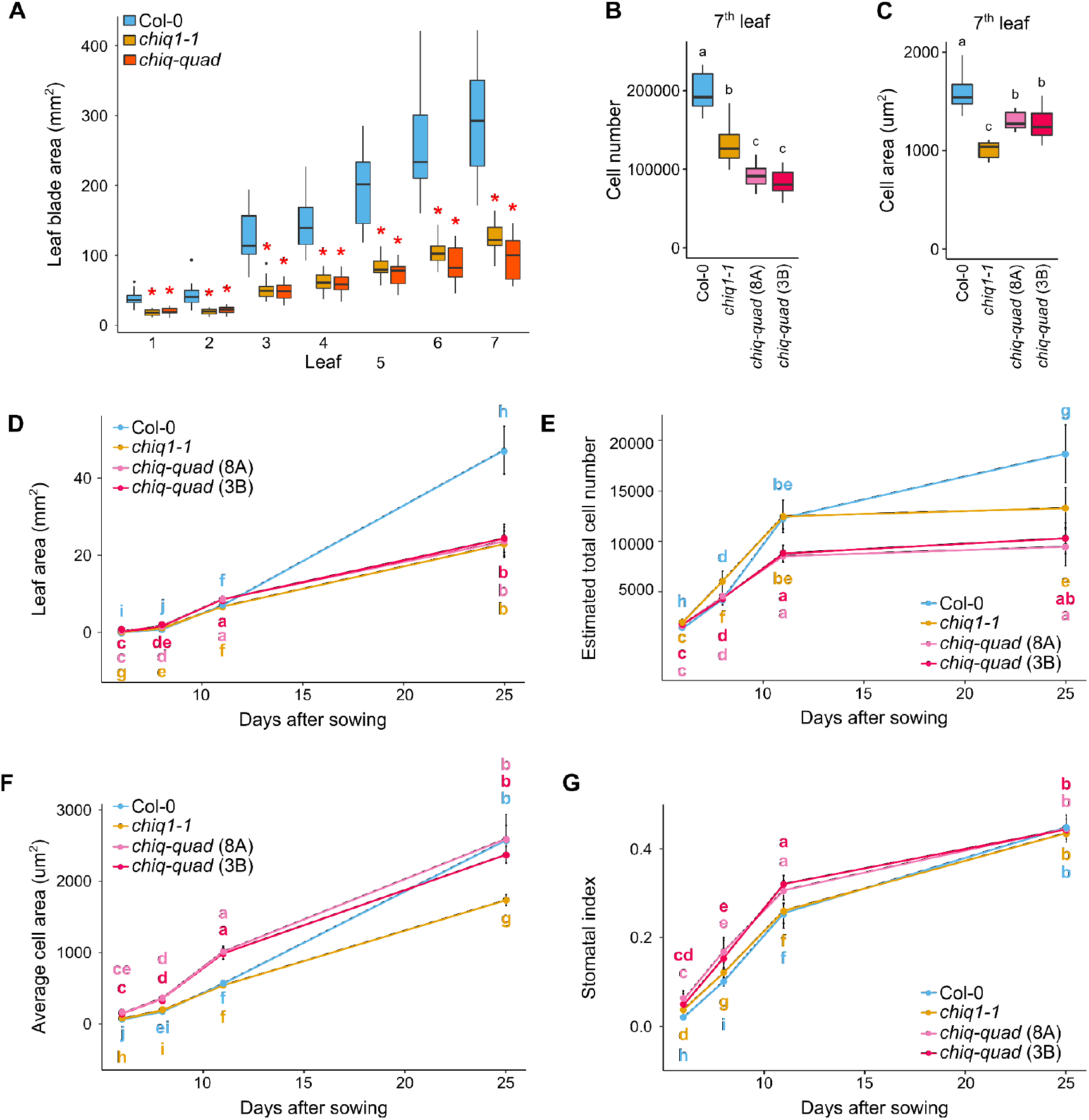
CHIQ proteins control the timing of the transition into differentiation. A) Leaf blade area (box and whisker plot) of mature leaves from soil-grown wild type (Col-0, blue), *chiq1-1* (orange), and a quadruple mutant (*chiq-quad: chiq1-1;chiql4;chiql5;chiql6*, red) (n = 17-22 per leaf per genotype; N = 3). The blade area of each leaf was compared among the three genotypes using Kruskal-Wallis test followed by pairwise comparison using the Wilcoxon test. The red asterisk indicates statistical significance (p-value < 0.05) compared to the wild type. B-C) Total cell number (B) and average cell size (C) in the abaxial epidermis of the 7^th^ leaf from 36 day-old wild type (Col-0, blue), *chiq1-1* (orange) and two quadruple lines carrying the plasma membrane marker RCI2A (*chiq-quad* (8A) (light pink) and *chiq-quad* (3B) (pink)) (n = 11 per genotype; N = 3). Letters represent significantly different groups (p-value < 0.05) as determined by two-way analysis of variance followed by post hoc Tukey’s test. D-G) Kinematic studies in the first pair of true leaves from wild type (Col-0, blue), *chiq1-1* (orange) and two quadruple lines carrying the plasma membrane marker RCI2A (*chiq-quad* (8A) (light pink) and *chiq-quad* (3B) (pink)) grown on soil. Leaf blade area (D), estimated total cell number (E), average cell area (F) and stomatal index (G) in the abaxial epidermis from seedlings at 6, 8, 11 and 25 days after sowing (n = 10-23 per genotype per time point; N = 3). Letters represent significantly different groups (p-value < 0.05) as determined using Kruskal-Wallis test followed by pairwise comparison using the Wilcoxon test. Graphs were generated using the ggplot2 package in R (49).

## Discussion

In this study, we show that CHIQ1 maintains the proliferative capacity of cells and delays the timing of cell cycle exit during organ development by combining cell population (kinematic analysis) and single cell (cell lineage tracking) studies. Kinematic studies consist of assessing cell size and cell number over time in growing organs and are a powerful framework to characterize how cellular parameters change during organ development at the organ level (33). However, during a kinematic study, proliferation rate and the length of the cell cycle are calculated from the estimated cell number at each time point assuming that all cells divide. This is not accurate based on how proliferation occurs in Arabidopsis leaves (25, 34). Nevertheless, twenty-one genes have been reported to affect proliferation rate according to kinematic studies (Table S1). Since cell cycle length was not measured directly in any of these studies, it is not possible to distinguish if these genes are involved in controlling the proportion of dividing cells or cell cycle length at this time.

Here, we calculated cell cycle length of wild type and *chiq1-1* leaves by tracking individual cells over time with live imaging and considering only the cells that underwent division. We found no difference in the cell cycle length between genotypes in both control conditions and in response to a G1/S cell cycle checkpoint trigger using HU. This discovery revealed that the proliferation rate at the organ level in *chiq1-1* was reduced due to the number of dividing cells decreasing rather than an increase in cell cycle length.

Duration of cell proliferation in an organ is controlled by both cell intrinsic (cell-level) and extrinsic (organ-level) mechanisms. How these mechanisms interact to control proliferation exit and final cell number in leaves is unknown. Several lines of evidence support that the duration of cell proliferation is controlled by cell extrinsic mechanisms that depend on cell-cell communication via the movement of proteins or small molecules (35–39). Among the three proposed cell-intrinsic mechanisms (counter, timer, sizer), the cell cycle counter mechanism has more supporting evidence than a timer or sizer mechanism in plants (19, 20). A cell cycle counter mechanism controlling stem cell division has been proposed for stem cells in floral and root meristems (19). The stem cell activity in the floral meristem terminates when the expression of the homeotic transcription factor *WUSCHEL* (*WUS*) is stably repressed by the transcriptional regulators *KNUCKLES* (*KNU*) and *AGAMOUS* (*AG*) (19). This repression depends on the dilution of the repressive chromatin mark H3K27me3 on the promoter of *KNU* across successive replication cycles, which allows for AG to increase *KNU* gene expression. This DNA replication-dependent delay in the activation of *KNU* expression provides the proper number of cells required to form floral organs by a cell division counter mechanism (19). Similarly, the amount of the chromatin mark H3K36me at specific target genes after each DNA replication round is controlled by the SET-domain protein ASH1-RELATED 3 (ASH1R3) (20). This epigenetic modification at each cell cycle could act as a cell division counter in the root apical meristem (20). In both cases, the dilution of a chromatin mark sets the timing of cell differentiation (and therefore may force cells to exit proliferation), indicating chromatin-level regulation. Although a similar mechanism could operate in leaves to control the number of cell division rounds within the amplifying cell population (i.e., dividing cells that are not stem cells), it would have to depend on other genes since *WUS* and *KNU* are stem-cell specific and *ASH1R3* is root specific, and none of these genes have been implicated in controlling the proliferation-differentiation transition in leaves. In this work, we showed that CHIQ1, and the CHIQ-like proteins CHIQL4, CHIQL5 and CHIQL6, control the proper timing of transition from proliferation into differentiation during organ development and, previously, we found that CHIQ1 interacted with CHIQL6, which interacted with a member of the chromatin repressor complex PRC2 (21).These results open the possibility that CHIQ-like proteins, including CHIQ1, might also control the number of times a cell divides by modulating the level of chromatin markers.

Most studies of cell proliferation and how different genes act to control proliferation rate have been performed at the cell population level using kinematic studie. While this approach is accurate in describing a general cellular phenotype (25, 40), it explains phenomena for an “average ideal cell” disregarding valuable information contained in the heterogeneous population. This study highlights the need to study development at the individual cell level in order to understand fundamental rules of biology such as what cell-intrinsic mechanisms control cell proliferation exit in plant organs.

## Material and Methods

### Plant material and growth conditions

*Arabidopsis thaliana* plants were grown either in PRO-MIX HP Mycorrhizae potting soil (Premier Tech Horticulture, Quakertown, PA) or in 0.5X Murashige and Skoog basal salt mixture (MS) media (PhytoTechnologies Laboratories) (pH 5.7), supplemented with 0.8% agar (Difco) and 1% sucrose (SIGMA). Plants were grown at 22°C in a 16:8 light:dark photoperiod, 40% RH, and ~115 μmol m^-2^s^-1^ measured at pot-level. Seeds were stratified at 4°C for four nights to break dormancy. The following SALK lines were used: SALK_064001 (*chiq1-1*), SALK_116702 (*chiql 4-1*), SALK_105421 (*chiql 5-1*) and SALK_086603 (*chiql 6-1*). The quadruple mutant was generated by crossing SALK_064001, SALK_11670, SALK_105421 and SALK_086603.

To construct a binary vector that expresses *CHIQ1*’s wild type allele, 642 bp of the promoter region (including the 5’ UTR) plus the coding region of AT2G45260 lacking the stop codon was amplified by PCR from Col-0 genomic DNA, cloned into the entry vector pDONR221 (Life Technologies), and transferred to the binary vector pGWB640 (41) using Gateway cloning (Life Technologies) to create the vector CHIQ1pro:CHIQ1-YFP.

To construct a binary vector that overexpresses *CHIQ1*’s wild type allele, the AT2G45260 (CHIQ1) protein-coding sequence was amplified by PCR from Col-0 genomic DNA, cloned into the entry vector pDONR221 (Life Technologies), and transferred to the binary vector pB7HFC3_0 (42), using Gateway cloning (Life Technologies) to create the vector 35Spro:CHIQ1-FLAG.

To obtain complemented lines, *chiq1-1* plants were transformed with the transgene *35Spro:CHIQ1-YFP* or *35Spro:CHIQ1-FLAG* by dipping their flowers into an Agrobacterium (strain pGVl101 pMP90) cell suspension. Briefly, a single colony of each construct was cultured in LB media with 50 ug/ml gentamicin, 25 ug/ml rifampicin and 50 ug/ml kanamycin or 50 ug/ml spectomicin until the culture reached an O.D. at 600 of 0.5-1. When the culture was ready, the LB was removed and the cell pellet was resuspended in plant growth media (0.43% MS (m/v), 0.05% MES (m/v), 5% sucrose (m/v), 0.02% Silwett L77 (v/v), pH 5.8-6) and transferred into a wide-mouthed container. Inflorescences of *chiq1-1* plants were dipped into this solution for a few minutes. Treated plants were placed horizontally and covered, left in the dark overnight, and moved to greenhouse-growing conditions the next day.

Transgenic lines were selected with BASTA (glufosinate ammonium). The adult phenotype was complemented in all transgenic lines analyzed (~10 independent lines per construct). The lines 3089x (CHIQ1pro:CHIQ1-YFP) and HFC10.4 (35Spro:CHIQ1-FLAG) were chosen for detailed organ size studies during development.

For marker gene studies, we used the following lines: 1) a transcriptional fusion containing the promoter of the cell cycle gene *CDKB1.1* linked to the GUS-encoding gene *uidA (CDKB1.1pro:GUS)* from Dr. Kathryn Barton (Carnegie Institution for Science, USA); 2) a transcriptional fusion containing the promoter of the cell cycle gene *CYCD3.3* linked to the GUS-encoding gene *uidA* (*CYCD3.3pro:GUS*) from Dr. Jose Dinneny (Stanford University, USA); 3) a fusion containing the promoter of the cell cycle gene *CYCB1.1* and its destruction box linked to the fluorescent protein GFP (*CYCBpro:CYCB_destruction box_*-GFP) from Dr. Masaaki Umeda (Nara Institute of Science and Technology, Japan); and 4) a translational fusion containing the promoter of the epidermis-specific gene *ATML1* and the plasma membrane localized protein RCI2A linked to the fluorescent protein mCitrine from Dr. Adrienne Roeder (Cornell University, USA). These transgenic lines were crossed with *chiq1-1* plants to obtain siblings carrying the corresponding marker gene in the wild type background or in the homozygous mutant background.

### Phenotypic analyses

#### Leaf area

Leaf area was measured from plants grown in soil for 4, 5, 6, 7, 8, 11, 13, 15, 19, 22 and 25 days after sowing. Leaves were dissected and photographed with a compound microscope (Nikon), dissecting scope (Leica MZ6 microscope), or scanned, depending on their size. Leaf blade area was measured from the images with ImageJ software (43).

#### Root length

Primary root length was measured in seedlings grown on 0.5X MS agar media supplemented with 1% sucrose for 3, 5, 7, 10, 12 and 14 days after stratification. Seedlings were imaged at the different time points and the length of the primary root was measured with ImageJ software (43). Root length in 21 day-old plants was measured using a rhizotron system (44). Assembled rhizotrons were placed in a box with water, and sown with seeds that had been stratified at 4°C for four days to break dormancy. These boxes were sealed with a transparent lid and packing tape. Plants were grown under a 16:8 light:dark photoperiod in a growth chamber at 22°C under full-spectrum LED lights outputting ~150 PPFD at pot level. Three days after sowing, the lids were removed, and the rhizotrons were watered once a day for seven days with 2 mL of distilled water using a micropipette. Twenty-one days after sowing, the rhizotrons were opened, and individual plants were removed, washed, and mounted onto a clear plastic sheet using double-sided tape. These plants were scanned and the root length of each plant was measured using ImageJ software (43).

#### Cell area and number in leaves

Cell area was measured from confocal images of the 1^st^, 4^th^ and 7^th^ leaves from 25-35 day-old plants (mature leaf) and the 1^st^ true leaves from 4, 6, 8, 11, 13, 15, 19, and 25 day-old plants (Col-0 and *chiq1-1*, growing leaf) or 6, 8, 11 and 25 day-old plants (quadruple mutant *chiq1-1;chiql4;chiql5;chiql6*, growing leaf) grown on soil. All lines (Col-0, *chiq1-1*, the complemented line 35Spro:CHIQ1-FLAG, and the quadruple mutant *chiq1-1;chiql4;chiql5;chiql6* express the plasma membrane fluorescent marker RCI2A in the epidermis (30).The fluorescent marker RCI2A was introgressed into *chiq1-1* and the 35Spro:CHIQ1-FLAG line; and introduced by Agrobacterium transformation into the quadruple background. To measure leaf area, leaves were dissected and photographed using a Leica SP8 confocal microscope, dissecting microscope (Leica MZ6 microscope) or scanner, depending on their size. Three confocal images (20x/0.7, glycerine, 512 x 512) of the abaxial epidermis were taken from each leaf, corresponding to the 25^th^, 50^th^, and 75^th^ percentile region of the blade. Excitation was performed using a 516-nm laser line, and mCitrine was collected at 526-580 nm. Maximum projections were calculated in ImageJ and all complete cells were traced (pencil width = 1 pixel for day 6 and day 8, pencil width = 2 pixels for day 11 and older plants). Total leaf area and individual cell areas were measured using ImageJ software (43). Average cell size, total cell number, cell production and stomatal index were calculated following (33).

#### Cell number in the root apical meristem (RAM)

Roots of 8 and 12 day-old wild type (Col-0) and *chiq1-1* seedlings were stained with 10 ug/ml propidium iodide (PI) for ~5 minutes and imaged with a Leica SP8 confocal microscope. Excitation was performed using a 488 nm laser line, and PI was collected at 570-670 nm. ImageJ software was used to count epidermal cells along a cell file. The RAM zone was defined as described in the literature (45): the region in the root encompassed by the first cell in the epidermal file after the quiescent center to the first cell that is at most half the length of the adjacent older cell within the same cell file.

### Cell cycle marker gene studies

Expression of the reporter gene *uidA* (GUS) driven by the promoter of the cell cycle gene *CYCD3.3 (CYCD3.3pro:GUS)* was analyzed in the 1^st^ true leaf of plants grown for 7, 8 and 9 days on 0.5X MS agar plates supplemented with 1% sucrose. Expression of GUS driven by the promoter of the cell cycle gene *CDKB1.1 (CDKB1.1pro:GUS)* was analyzed in the 4^th^ and 5^th^ leaf of soil-grown plants at 16, 18, 20, 22 and 24 days after sowing (DAS). Plants were stained in GUS staining solution (46) at 37°C overnight, and were destained in 70% ethanol at room temperature for 24 hours. Images were taken with a Leica MZ6 microscope. Expression of GFP driven by the promoter and destruction box of the cell cycle gene *CYCB1.1* (*CYCBpro:CYCB_destruction box_*-GFP) was analyzed in the root apical meristem of 5 day-old seedlings grown on plates. Roots were stained in 10 ug/ml propidium iodide for ~5 minutes before imaging with a Leica SP8 confocal microscope. Excitation was performed using a 488-nm laser line, and PI and GFP were collected at 570-670 and 498-548 nm, respectively.

### Endoreduplication analysis

Mature 6^th^ and 7^th^ leaves were collected from plants grown in soil. Leaves were chopped into thin slices and incubated in an enzyme solution for 90 minutes to obtain protoplasts (47). Protoplasts were filtered with a 100 um cell strainer and washed several times with W5 buffer (47). After the last wash, protoplasts were resuspended in lysis buffer (45mM magnesium chloride, 30mM sodium citrate, 20mM MOPS, 0.5% triton, 2% 4’,6-diamidino-2-phenylindole (DAPI, Thermo Scientific), pH 7) (48) to obtain nuclei. Nuclei solution was filtered with a 40um cell strainer before flow cytometry analysis. Data were collected on the LSR II.UV analyzer (NIH S10 Shared Instrument Grant S10RR027431-01) in the Stanford Shared FACS Facility (SSFF) and analyzed using the flowCore, flowViz, flowStats, and flowDensity libraries within the R package Bioconductor (https://www.bioconductor.org/).

### Analysis of cell cycle parameters

Six-day old wild type and *chiq1-1* seedlings carrying a plasma membrane fluorescent marker RCI2A in the epidermis (30) and grown on agar media were mounted on slides with 400ul of 0.5X MS media with 1% sucrose, after removal of one cotyledon. For HU treatments, seedlings were mounted on slides with 400ul of 0.5X MS media with 1% sucrose plus 0.5mM or 1mM HU. Exposed young leaves were imaged using a Leica SP8 confocal microscope through a multi-immersion 20x/0.7 objective using glycerine immersion media. Excitation was performed at 516 nm, and mCitrine was collected at 526-580 nm. Leaves were imaged taking a z-stack once a day for 2 days. Maximum projections were calculated in ImageJ (43).

The mid-region of the leaf, where active division is happening, was determined and an area of approximately 40 cells was selected. Cells within the selected region were traced (pencil width = 1 pixel) and cell sizes were measured using ImageJ software (43). To measure cell division, the cells within the region were numbered and colored using ImageJ (43). The mother and daughter cells were manually tracked through subsequent images. Cells that divided were identified and their progeny counted. The number of cells produced per mother cell was used to calculate proliferation rate and cell cycle length.

## Acknowledgements

We thank ABRC and Drs. K. Barton, M. Umeda, A. Roeder, and J. Dinneny for providing mutant seeds and seeds of cell cycle and plasma membrane marker lines; D. Nusinow and S. Ishiguro for providing binary plasmids; T. LaRue, A. Srinivas, and Dr. J. Vilarasa-Blasi for technical assistance on the rhizotron and root imaging studies; H. Nam and J. A. Kim for technical assistance; T Knaak and the SSFF for technical assistance on endoreduplication studies; A. Malkovskiy for microscopy assistance, G. Materassi-Shultz for maintenance and watering all plants used in this study, and Drs. D. Ehrhardt, H. Meyer, S. Xu, L. de Veylder, L. Willis, and members of the Rhee lab and the Carnegie-Stanford plant research community for helpful discussion. This work was done on the ancestral land of the Muwekma Ohlone Tribe, which was and continues to be of great importance to the Ohlone people.

## Authors’ contributions

F.B. and S.Y.R. conceived the project, F.B conducted the mature plant and leaf phenotypic characterization of *chiq1-1*, complemented lines and the quadruple mutant, B.J. performed root studies in *chiq1-1* and complemented lines, F.B. and B.J. generated the quadruple mutant and performed cell cycle marker studies in leaves and roots, F.B. performed cell lineage tracking experiments, F.B., E.L. and S.Y.R. processed and analyzed imaging data from cell lineage tracking experiments, F.B. and E.L. processed and analyzed the cellular phenotype of *chiq1-1*, complemented lines and the quadruple mutant lines, H.C. assisted with confocal microscopy and image analysis. Y.D. provided intellectual contribution and input in the manuscript organization, F.B. wrote the manuscript, and B.J., Y.D. H.C., and S.Y.R. edited the manuscript.

## Competing interests

Authors declare no conflict of interest.

## Funding

This work was supported in part by Carnegie Institution for Science Endowment and grants from the National Science Foundation (IOS-1546838, IOS-1026003) and the U.S. Department of Energy, Office of Science, Office of Biological and Environmental Research, Genomic Science Program grant nos. DE-SC0018277, DE-SC0008769, DE-SC0020366, and DE-SC0021286.

## Data availability

Data is available in the main text or the supplementary materials.

## Authors’ information

Correspondence and requests for materials should be addressed to S.Y.R.

## Supplementary Information

## Notes

### Competing Interest Statement

The authors have declared no competing interest.

### Summary of Updates

Introduction, Figs 4A-B and Fig S4 and S5, and References updated. More data points (experiments) were added to Fig 4A-B and Fig S4 and S5. A sentence was added to the introduction and a reference describing CHIQ1's role in stress was included.

