## Supplemental information for "*CHIQUITA1* maintains temporal transition between proliferation and differentiation in *Arabidopsis thaliana*"

### Supplementary Information

File contains 7 figures and 1 table

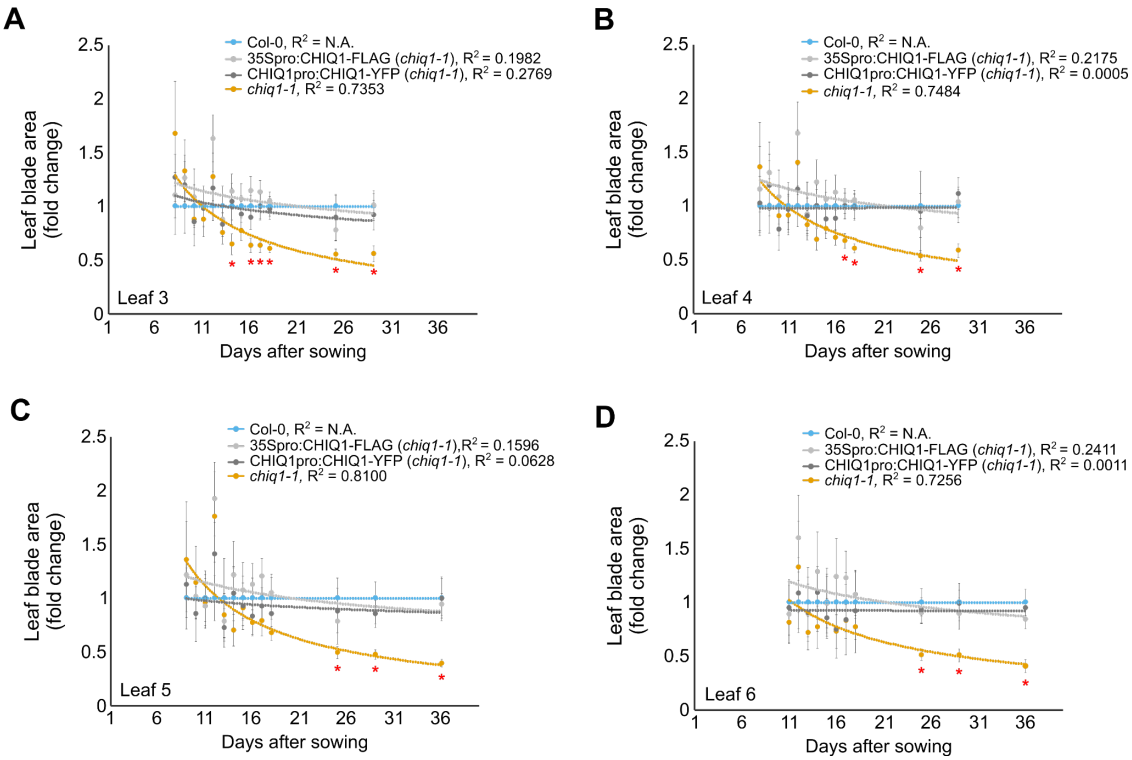


Figure S1

***chiq1-1* young leaves are the same size as in wild type.**

A-D) Scatter plot of normalized leaf blade area against the wild type value of the same age of the third (A), forth (B), fifth (C), and sixth (D) leaf in soil-grown plants ranging in age from 4 days to 36 days from wild type (blue), *chiq1-1* (orange), and two complemented lines (*CHIQ1pro:CHIQ1-YFP* line, dark gray and *35Spro:CHIQ1-FLAG*, light gray) (n = 6-10 per leaf per genotype; N = 1). Error bar represents 95% confidence interval. The regression line corresponds to a power regression. All genotypes were compared at each time point using two-way analysis of variance followed by post hoc Tukey's test (p-value < 0.05). Red asterisks represent a significant difference between *chiq1-1* and the other three genotypes

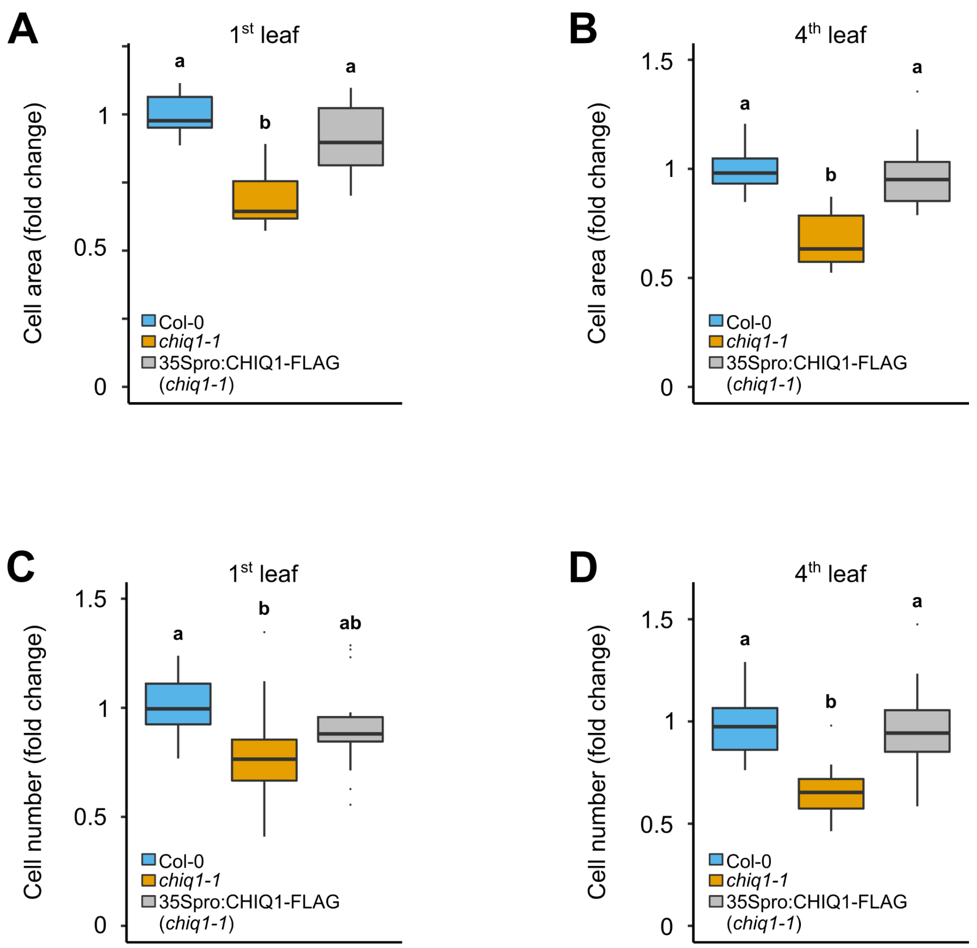


Figure S2

***chiq1-1* mature leaves have fewer and smaller cells.**

A-B) Normalized average cell size in the abaxial epidermis of the 1st (A) and 4th (B) leaf against the wild type value of the same experiment from wild type (Col-0, blue), *chiq1-1* (orange) and a complemented line (*35Spro:CHIQ1-FLAG*, gray) (n= 9-24 per genotype; N=4). C-D) Normalized total cell number in the abaxial epidermis of the 1st (C) and 4th (D) leaf against the wild type value of the same experiment from wild type (Col-0, blue), *chiq1-1* (orange) and a complemented line (*35Spro:CHIQ1-FLAG*, gray) (n= 9-24 per genotype; N=4). Letters represent significantly different groups (p-value < 0.05) as determined by two-way analysis of variance followed by post hoc Tukey's test.

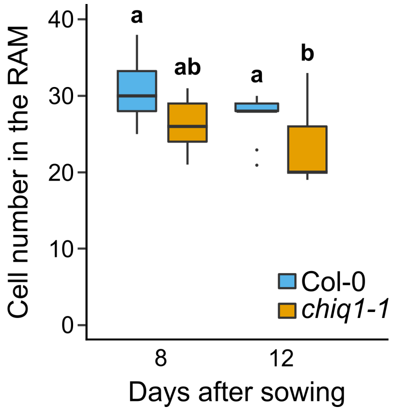


Figure S3

**The number of epidermis cells in the root apical meristem (RAM) is smaller in *chiq1-1* than in wild type**

Number of cells in the epidermis of the RAM of 8 and 12 day-old wild type (blue) and *chiq1-1* (orange) seedlings (n = 10-13 per time point per genotype; N=3). Letters represent significantly different groups (p-value < 0.05) as determined by two-way analysis of variance followed by post hoc Tukey's test.

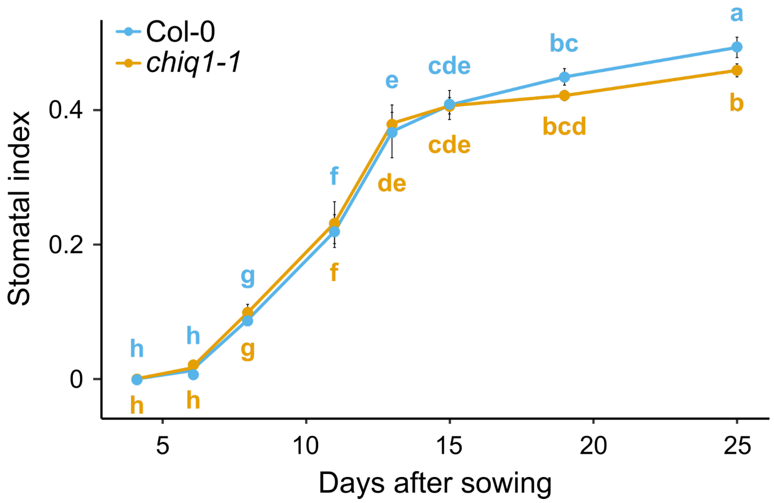


Figure S4

**Stomatal index in wild type and *chiq1-1* seedlings during development**

Stomatal index in the abaxial epidermis from wild type (Col-0, blue) and *chiq1-1* (orange) seedlings at 4, 6, 8, 11, 13, 15, 19 and 25 days after sowing (n = 5-30 per genotype per time point; N = 1-4). Letters represent significantly different groups (p-value < 0.05) as determined by two-way analysis of variance followed by post hoc Tukey's test.

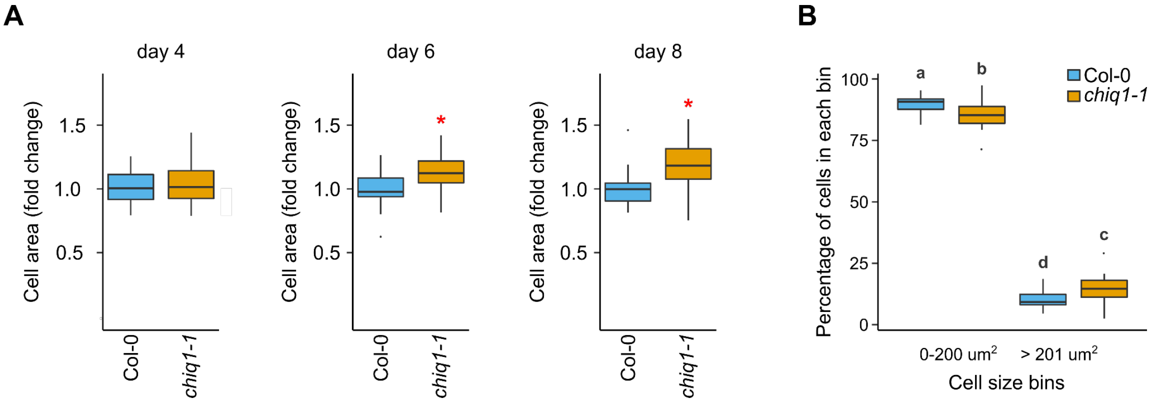


Figure S5

**Average cell area is greater in *chiq1-1* young leaves than in wild type and the number of dividing cells is smaller in *chiq1-1* 8-day old leaves than in wild type**

A) Normalized average cell area against the wild type value for each experiment from 4 (left), 6 (middle) and 8-day old (right) first true leaves from wild type (Col-0, blue) and *chiq1-1* (orange) seedlings (n = 20-30 per genotype per time point, N = 3-4). These graphs were generated with the same data as in Fig. 4A. Red asterisks represent significantly different groups (p-value < 0.05) as determined by Student t-test. B) Percentage of cells (excluding guard cells) within a particular cell size range in 8 day-old wild type and *chiq1-1* leaves (n = 20 per genotype, N = 4). Letters represent significantly different groups (p-value < 0.05) as determined by two-way analysis of variance followed by post hoc Tukey's test.

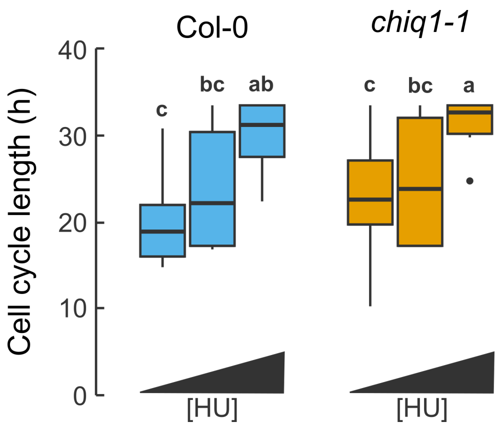


Figure S6

**Hydroxyurea (HU) treatment delays the cell cycle in wild type and *chiq1-1* cells**

Cell cycle length in wild type (Col-0, blue) and *chiq1-1* (orange) seedlings growing with 0mM, 0.5mM or 1mM HU (n = 6-15 per genotype per treatment; N = 2-3). Letters represent significantly different groups (p-value < 0.05) as determined by two-way analysis of variance followed by post hoc Tukey's test. Graphs were generated using the ggplot2 package in R (49).

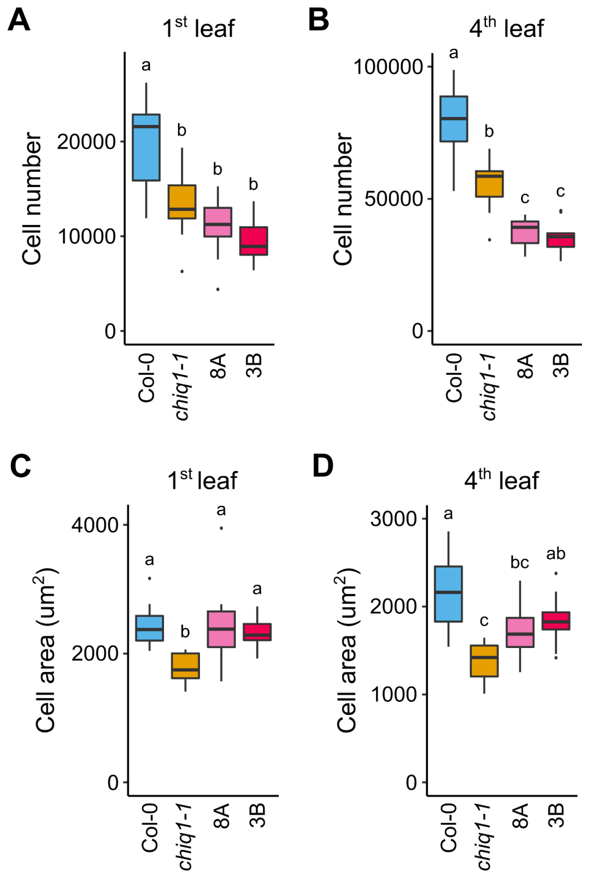


Figure S7

**CHIQ proteins decrease final cell number in leaves**

A-B) Total cell number in the abaxial epidermis of the 1st (A) and 4th (B) leaf from wild type (Col-0, blue), *chiq1-1* (orange) and two quadruple mutants carrying the plasma membrane marker RCI2A (*chiq-quad* (8A) (light pink) and *chiq-quad* (3B) (pink)) (n = 10-11 per genotype; N = 3). C-D) Average cell size in the abaxial epidermis of of the 1st (C) and 4th (D) leaf from wild type (Col-0, blue), *chiq1-1* (orange) and two quadruple mutants carrying the plasma membrane marker RCI2A (*chiq-quad* (8A) (light pink) and *chiq-quad* (3B) (pink)) (n = 10-11 per genotype; N = 3). Letters represent significantly different groups (p-value < 0.05) as determined by two-way analysis of variance followed by post hoc Tukey's test. Graphs were generated using the ggplot2 package in R (49).

Table S1

List of paper reporting kinematic analysis in determinate organs (leaves, petals) of *Arabidopsis thaliana*

OX: overexpression line

| Paper | System | Gene | Molecular function | Is proliferation rate affected? | Is the duration of proliferation affected? | Is cell cycle length affected? |
|---|---|---|---|---|---|---|
| Functional Analysis of Cyclin-Dependent Kinase Inhibitors of Arabidopsis | leaf 1 and 2 (OX) | KRP2 | CDK-inhibitor | Yes | No | No (but measured from kinematic) |
| Multiple mechanisms explain how reduced KRP expression increases leaf size of Arabidopsis thaliana | leaf 1 and 2 (mutant) | KRP4/6/7 | CDK-inhibitor | No | Yes | No (but measured from kinematic) |
| Control of Plant Organ Size by KLUH/CYP78A5-Dependent Intercellular Signaling | petal (mutant) | KLUH/CYP78A5 | Cytochrome | Yes | Yes | Not measured |
| Cell number and leaf development in Arabidopsis: A functional analysis of the STRUWWELPETER gene | leaf 1 and 2 (mutant) | SWP/MED 14 | Mediator subunit | Yes | Yes | No (but measured from kinematic) |
| The Arabidopsis thaliana NGATHA transcription factors negatively regulate cell proliferation of lateral organs | leaf 1 and 2 (OX) | NGA1 | TF (RAV subfamily of the B3-type TF superfamily) | Yes | No | Not measured |
| | | NGA4 | TF (RAV subfamily of the B3-type TF superfamily) | Yes | Yes | Not measured |
| | leaf 1 and 2 (mutant) | NGA1, NGA2, NGA3, NGA4 | TF (RAV subfamily of the B3-type TF superfamily) | Yes | No | Not measured |
| SAMBA, a plant-specific anaphase-promoting complex/cyclosome regulator is involved in early development and A-type cyclin stabilization | leaf 1 and 2 (mutant) | SAMBA | APC/C regulator | No | No | Not measured |
| GROWTH-REGULATING FACTOR 9 negatively regulates arabidopsis leaf growth by controlling ORG3 and restricting cell proliferation in leaf primordia | leaf 1 and 2 (mutant and OX) | GRF9 | TF | Yes | No | Not measured |

| | | | | | | |
|---|---|---|---|---|---|---|
| Translationally controlled tumor protein is a conserved mitotic growth integrator in animals and plants | leaf 1 and 2 (RNAi) | TCTP | | Yes | No | Yes (but measured from kinematic) |
| ORESARA15, a PLATZ transcription factor, mediates leaf growth and senescence in Arabidopsis | leaf 1 and 2 (mutant) | ORE15 | PLATZ TF | Yes | Yes | Not measured |
| The DNA replication checkpoint aids survival of plants deficient in the novel replisome factor ETG1 | leaf 1 and 2 (mutant) | ETG1 | replisome factor | Yes | No | Yes (but measured from kinematic)/ FC data supports cell cycle length defects) |
| A Membrane-Bound NAC Transcription Factor Regulates Cell Division in Arabidopsis | leaf 1 and 2 (mutant) | NTM1 | NAC TF | Yes | Yes | Not measured |
| Histidine Kinase Homologs That Act as Cytokinin Receptors Possess Overlapping Functions in the Regulation of Shoot and Root Growth in Arabidopsis | leaf 1 and 2 (mutant, double and triple) | AHK | Kinase (two component system) | Yes | Maybe? | Not measured |
| Enhanced cytokinin degradation in leaf primordia of transgenic Arabidopsis plants reduces leaf size and shoot organ primordia formation | leaf 1 and 2 (OX) | CKX3 | cytokinin oxidase/dehydrogenase | Yes | Yes | Not measured |
| The E3 Ubiquitin Ligase BIG BROTHER Controls Arabidopsis Organ Size in a Dosage-Dependent Manner | petal (mutant, OX) | BB | E3 ligase | Not measured | Yes | Not measured |
| Control of final seed and organ size by the DA1 gene family in Arabidopsis thaliana | petal (mutant) | DA1 | ubiquitin receptor | Not measured | Yes | Not measured. |
| ROTUNDIFOLIA4 Regulates Cell Proliferation Along the Body Axis in Arabidopsis Shoot | leaf 1 and 2 (mutant) | ROT4 | small protein (6.2 kDa) | Yes | No | Not measured |
| The Arabidopsis GRF-INTERACTING FACTOR Gene Family Performs an Overlapping Function in Determining Organ Size as Well as Multiple Developmental | leaf 1 and 2 (mutant) | GIF1/2/3 | TR | Yes | Yes | Not measured |

| | | | | | | |
|---|---|---|---|---|---|---|
| Properties | | | | | | |
| Analysis of Leaf Development in fugu Mutants of Arabidopsis Reveals Three Compensation Modes That Modulate Cell Expansion in Determinate Organs | leaf 1 and 2 (mutant) | FUGU2 | | Yes | No | Not measured |
| | | FUGU5 | | Yes | No | Not measured |
| | | ERECTA | Receptor-like kinase | Yes | No | Not measured |
| | | KRP2 | CDK-inhibitor | Yes | Yes | Not measured |
| | | AN3 | TR | No | Yes | Not measured |
| The Arabidopsis thaliana Homolog of Yeast BRE1 Has a Function in Cell Cycle Regulation during Early Leaf and Root Growth[ | leaf 1 and 2 (mutant) | HUB1 | RING E3 ligase | Yes | No | Yes (but measured from kinematic) |
| Arabidopsis SMALL ORGAN 4, a homolog of yeast NOP53, regulates cell proliferation rate during organ growth | leaf 5 (mutant) | SMO1 | nucleolar protein (NOP53 homolog) | Yes | No | Not measured |
